# scDIG: An R Shiny Application for Interactive Density-Based Gating of Single-Cell Proteomic and Transcriptomic Data

**DOI:** 10.64898/2026.05.20.726609

**Authors:** Polina Bombina, Anusha Bellapu, Lauren Fogel, Kevin Coombes, Klaus Ley

## Abstract

Delineating biologically meaningful cell populations within single-cell embedding spaces requires methods that balance expert guidance with reproducibility. We present **scDIG**, a Shiny-based tool that integrates bimodal index-driven feature selection, feature-weighted kernel density estimation, and interactive contour-based gating to define cell populations directly within two-dimensional projections of scRNA-seq and CITE-seq data. We applied **scDIG** to CITE-seq PBMC data from human subjects in the Cardiovascular Assessment Virginia (CAVA) cohort and show that it resolves transcriptionally distinct CD4^+^ T cell subpopulations within continuous embeddings that are not readily captured by conventional clustering approaches. These findings demonstrate the utility of **scDIG** for robust, reproducible classification of single-cell populations and for identifying immunologically relevant effector states. The app is freely available for non-commercial use at https://au-cbgm-shiny.augusta.edu/gating, with source code available at https://gitlab.com/pbombina/scdig.

## Introduction

Single-cell RNA sequencing (scRNA-seq) has revolutionized our understanding of cellular heterogeneity by enabling transcriptomic profiling of individual cells at unprecedented resolution. A fundamental challenge in scRNA-seq analysis is the identification and delineation of biologically distinct cell populations within low-dimensional embedding spaces produced by dimensionality reduction techniques such as UMAP^1^ and t-SNE^2^.

In flow cytometry, “gating” provides a systematic framework for identifying cell populations through sequential selection based on measurable cellular properties^3^. This paradigm combines the interpretability of manual boundaries with standardized, reproducible workflows. While scRNA-seq analysis would benefit from an analogous approach, current methods occupy two extremes. Manual selection of cell populations in embedding space, though allowing integration of expert knowledge, suffers from high inter-operator variability and poor reproducibility^4^. At the other extreme, fully automated graph-based clustering algorithms like Louvain^5^ or Leiden^6^ remove researcher control and frequently merge rare but biologically significant populations into larger clusters, obscuring important cellular subsets.

This gap motivates the need for semi-automated approaches that preserve expert guidance while providing objective, reproducible population boundaries. Here, we present **scDIG**, a Shiny-based tool that applies density-based gating to scRNA-seq embedding spaces, bridging the interpretability of manual gating with the rigor of algorithmic cell population definition.

## Methods

Figure 1 contains a summarized overview of our software implementation.

**Figure 1:**
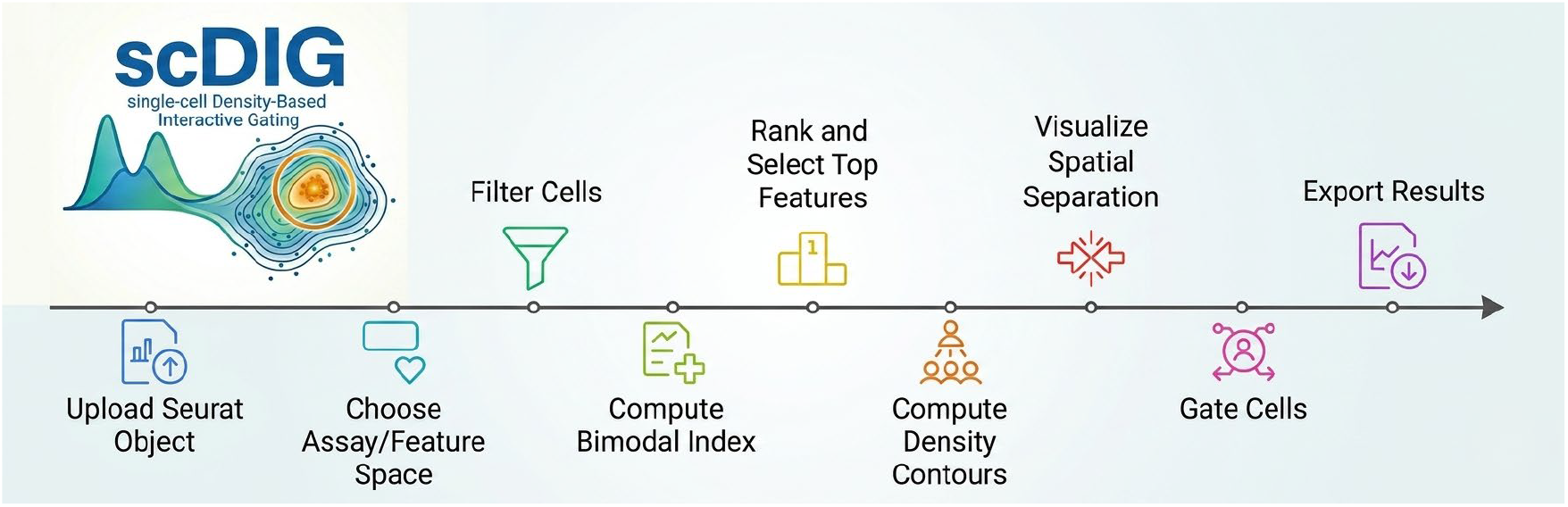
Steps of the analytical pipeline that is internally implemented in scDIG.

Users upload preprocessed data as Seurat objects in rds format. Upon loading, the application validates that the uploaded file is a Seurat object and extracts available assay names, dimensionality reductions, and metadata columns. Users then select the assay and dimensional reduction to use for downstream analysis. The UMAP reduction is selected by default if available.

### Cell Subsetting and Filtering

The application provides interactive cell subsetting capabilities through three options: selection of all cells, filtering by a single metadata column, or application of multi-column filtering criteria.

### Bimodal Index Calculation for Feature Selection

To identify features (genes or surface markers) exhibiting bimodal expression patterns, we computed the Bimodal Index (BI) for the selected cell subset^7^ using `bimodalIndex` function from the **BimodalIndex** package. Users can optionally restrict the analysis to variable features identified by Seurat’s `VariableFeatures()` function. To reduce computational burden and remove noise in scRNA-seq data, we applied pre-filtering before BI calculation. Genes expressed in fewer than 1% of cells (minimum 3 cells) were excluded, as were genes with mean expression below 0.01 (units). This filtering step removes likely technical noise while preserving biologically relevant features.

The BI calculation employs an adaptive sampling strategy that balances computational efficiency with statistical robustness based on dataset size:

- **Small datasets (≤15**,**000 cells)**: All cells are used in a single BI calculation
- **Medium datasets (15**,**001–30**,**000 cells)**: 80% of cells are randomly sampled across multiple iterations
- **Large datasets (>30**,**000 cells)**: 25,000 cells are randomly sampled across multiple iterations

For datasets requiring sampling, the calculation is repeated across three independent iterations with different random cell subsets. Within each iteration, genes lacking expression variance in the sampled cells are removed prior to BI computation. Final BI values represent the mean across all successful iterations.

If some iterations fail due to computational issues, the application issues a warning and computes the mean from available iterations.

Features are selected using either a user-specified BI threshold or a percentile-based cutoff (90th, 95th, or 99th percentile) derived from the BI distribution. Percentile-based selection offers a data-adaptive alternative by defining thresholds relative to the observed BI distribution within each dataset. Instead of relying on an absolute BI value, features are ranked by BI and only the most extreme fraction is retained. This approach accounts for dataset-specific differences in BI scale and variability, enabling consistent identification of highly bimodal and biologically informative features even when absolute BI values are not directly comparable across datasets. Results are presented in an interactive table, and histogram visualizations display BI distributions with threshold demarcation lines to guide transparent feature selection.

### Feature-Weighted Kernel Density Estimation

To visualize the spatial distribution of feature-enriched cell populations within two-dimensional embedding spaces, we implemented a feature-weighted kernel density estimation (KDE) approach. Cells can be colored by any high-BI feature using customizable color scales. This KDE method identifies regions where cells with elevated expression of selected features are concentrated, revealing relationships between molecular signatures and embedding topology.

Two-dimensional KDE is computed using bivariate Gaussian kernels via the kde2d function from the **MASS** package. Contour lines representing iso-density regions are extracted from the estimated density surface using the `contourLines` function (**grDevices** package). Each contour corresponds to a level set indicating regions of equivalent feature-weighted density. Users can interactively adjust the minimum contour threshold to focus on high-density, feature-enriched regions.

### Interactive Gating and Cell Selection

Users perform gating by clicking on the contours of interest in a two-dimensional plot. Gates are stored as named lists mapping gate identifiers to vectors of cell barcodes. Each saved gate preserves the complete set of cell identifiers rather than coordinate information, ensuring gates remain valid across different visualizations and data subsets. Gate names are automatically incremented using timestamps (format: `Gate_HHMMSS`) to prevent accidental overwrites.

During selection, real-time statistics display: - Number and percentage of selected cells relative to the current view - Sample cell barcodes (up to 5) for verification - Validation warnings for invalid selections

For contour-based analysis, cells falling within all displayed contour boundaries are automatically identified and quantified, providing objective assessment of feature-enriched regions.

### Gate Management and Analysis

The application supports several gate manipulation operations:

- **Copy**: Duplicate existing gates with new identifiers for creating variations or backups
- **Clear All**: Remove all gates simultaneously with user confirmation
- **Export/Import**: Save and load gates as rds files, including metadata (subset description, timestamp, subset cell identifiers)

Gate statistics are computed and visualized in a dedicated analysis panel:

#### Population Distribution

Bar plots display absolute cell counts and relative percentages for each gate. Gates are ordered by decreasing size with text annotations showing exact counts and percentages.

#### Overlap Analysis

A matrix quantifies pairwise cell overlap between all gates, enabling assessment of population relationships.

#### Subset Stability

When analyzing filtered subsets, the application calculates the percentage of gated cells retained in the current filtered view, allowing evaluation of gate stability across different data partitions.

Screenshots illustrating each step of the application workflow are provided in the Supplementary Material.

### Deployment and Availability

The application was developed using the R programming language (version 4.5.3) and the Shiny framework. It is deployed on a Linux-based server running Red Hat Enterprise Linux (RHEL) 8.10 and is publicly accessible at https://au-cbgm-shiny.augusta.edu/gating/ without user authentication. A reverse proxy (NGINX) is used to manage incoming traffic, with secure communication enabled via SSL/TLS encryption.

The application supports multiple concurrent users; however, due to the single-threaded nature of the open-source Shiny framework, computationally intensive tasks may temporarily impact responsiveness for other users. Source code for the application is available at https://gitlab.com/pbombina/scdig.

## Case study

To demonstrate the biological utility of our gating framework, we applied the app to CITE-seq data from PBMCs from subjects in the CAVA cohort (n = 61 patients and 51 markers)^8^, using the thresholded antibody-derived tag (ADT) assay for surface protein quantification. The full PBMC embedding encompassed all major immune cell populations, including CD4^+^ T cells (CD4T), CD8^+^ T cells (CD8T), NK cells, B cells, classical monocytes (cMo), intermediate monocytes (iMo), and non-classical monocytes (nMo), as visualized in the UMAP embedding (Figure 2A).

**Figure 2.**
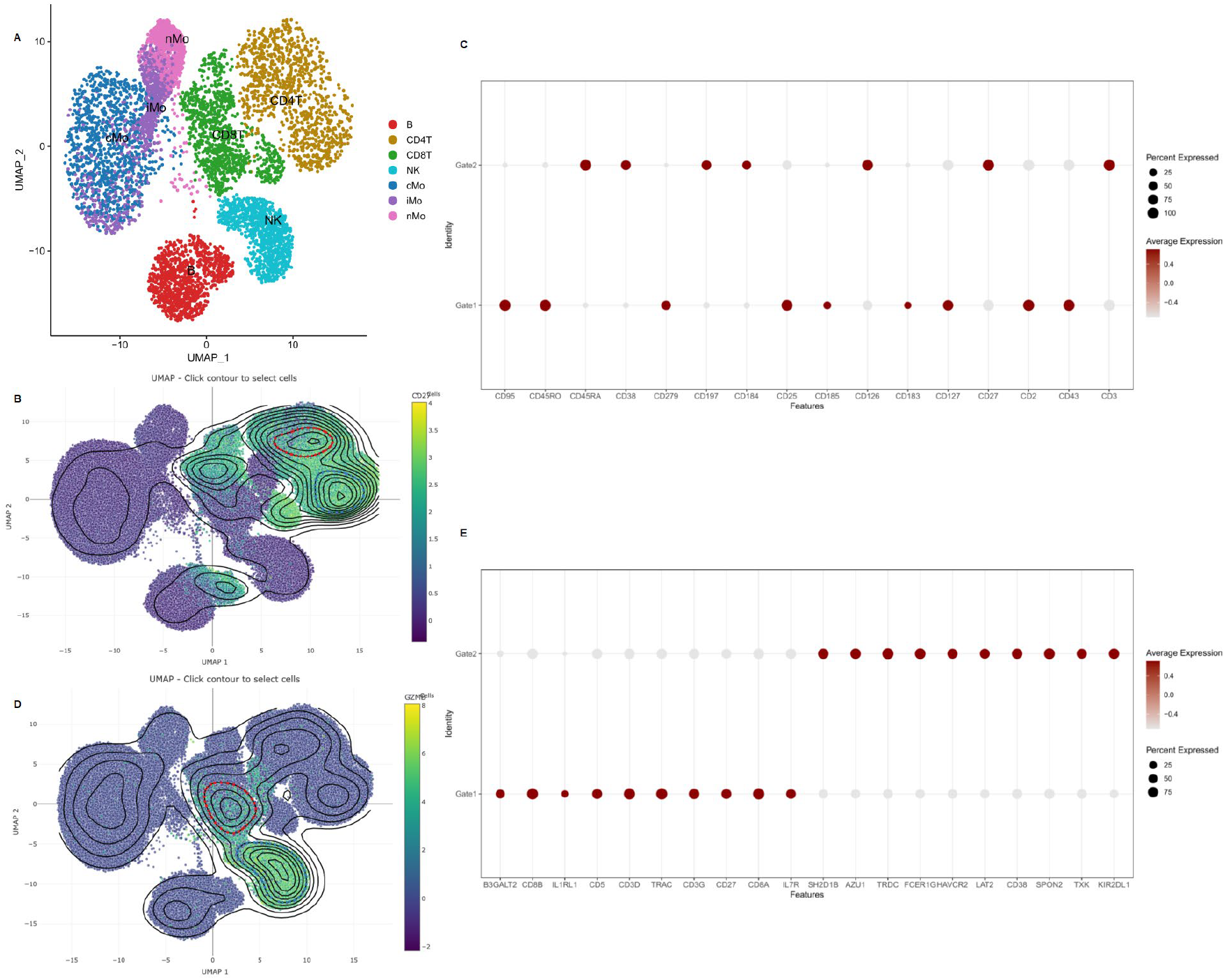
Density-based gating of cells from the CAVA cohort. (A) UMAP of 118,275 CITE-seq cells from 61 patients, colored by cell type. (B) Feature-weighted KDE of CD27 expression in CD4^+^ T cells; red and blue contours indicate Gate 1 and Gate 2 boundaries, respectively. (C) MAST differential ADT expression between Gate 1 and Gate 2, revealing distinct effector memory (Gate 1) and naïve/central memory (Gate 2) immunophenotypes. (D) Feature-weighted KDE of GZMB RNA expression; contours identify two transcriptionally defined high-density regions within the cytotoxic lymphocyte compartment, corresponding to Gate 1 and Gate 2. (E) MAST differential gene expression analysis between Gate 1 and Gate 2, revealing distinct conventional CD8^+^ T cell (Gate 1) and NK/innate-like lymphocyte (Gate 2) transcriptional phenotypes.

**Figure 3:**
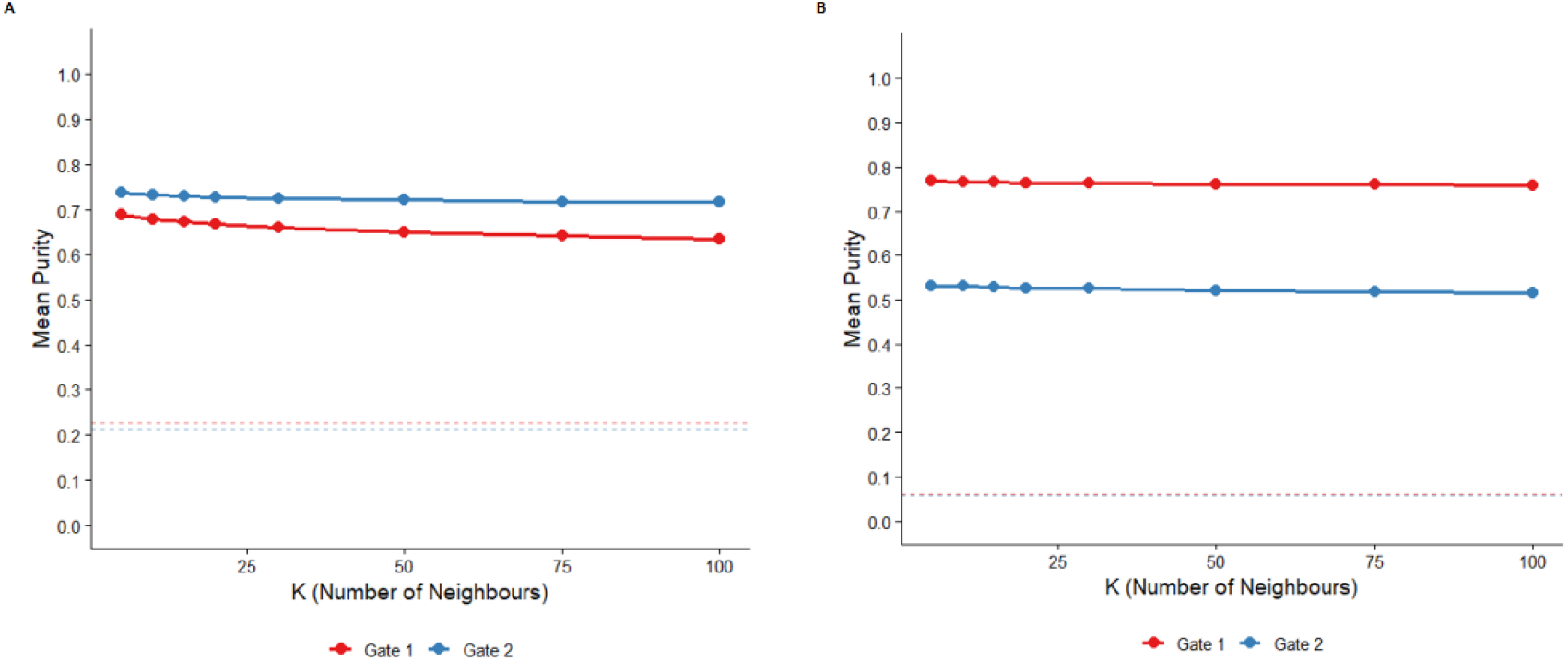
Sensitivity analysis of neighborhood purity. Mean nearest-neighbor (mNN) purity scores were calculated across neighborhood sizes (K = 5 to 100). Solid lines represent the observed purity for Gate 1 and Gate 2. Dashed lines indicate the random baseline. **(A)** mNN purity for gates defined using CD27 expression in CD4^+^ T cells from the ADT assay. **(B)** mNN purity for gates defined using GZMB expression from the RNA assay.

CD27, a surface marker exhibiting high expression variability across the CD4^+^ T cell embedding, was selected for feature-weighted kernel density estimation. Overlaying KDE contours onto the UMAP revealed two spatially distinct high-density regions, reflecting transcriptional heterogeneity not captured by standard clustering algorithms. Contour-based gating of this density peaks defined two populations: Gate 1 (9,201 cells) and Gate 2 (8,711 cells). Differential expression between gates was assessed using MAST^9^, a hurdle model designed for single-cell data that accounts for zero-inflation and expression variability. Gate 1 was enriched for CD95, CD45RO, PD-1 (CD279), CD25, CXCR5 (CD185), and CXCR3 (CD183), consistent with an effector and activated memory T cell phenotype. Elevated PD-1 expression is consistent with chronic antigen stimulation in the context of cardiovascular disease, while CXCR5 co-expression is suggestive of a T follicular helper (Tfh)-like subset. Gate 2 was enriched for CD45RA, CCR7 (CD197), CD38, and CXCR4 (CD184), reflecting a naïve or central memory phenotype with preserved lymph node homing capacity. This example demonstrates how contour-based gating can resolve biologically meaningful CD4+ T cell subsets from a continuous embedding, capturing immunologically relevant effector states that would otherwise remain indistinguishable within a single annotated cluster.

To demonstrate the applicability of **scDIG** to gene expression data, we applied the tool to the RNA assay of the same CITE-seq dataset from the CAVA cohort, using transcriptomic profiles rather than surface protein markers. Bimodal Index-based feature selection identified GZMB (Granzyme B) a serine protease central to cytotoxic effector function as a top-ranked gene exhibiting bimodal expression across the full PBMC embedding. GZMB was therefore selected for feature-weighted kernel density estimation. Overlaying KDE contours onto the UMAP colored by GZMB RNA expression revealed two spatially distinct high-density regions enriched for GZMB signal within the cytotoxic lymphocyte area of the embedding, and contour-based gating defined two populations: Gate 1 (8,430 cells) and Gate 2 (7,993 cells) (Figure 2D). Differential expression was similarly assessed using MAST. Gate 1 was enriched for canonical CD8^+^ T cell markers including CD8B, CD3D, TRAC, CD3G, CD27, CD8A, and IL7R, consistent with a conventional αβ T cell identity. Gate 2 showed high expression of SH2D1B, AZU1, TRDC, FCER1G, HAVCR2, LAT2, CD38, SPON2, TXK, and KIR2DL1, a signature associated with NK cells and innate-like lymphocytes including γδ T and NKT subsets (Figure 2E).

To assess the structural stability of the gated populations, we performed a sensitivity analysis on the neighborhood parameter (*K*). The mean nearest-neighbor purity was calculated iteratively across a range of values (*K* ∈ {5,10,15,20,30,50,75,100}). This allowed us to observe how local versus global neighborhood definitions affected the coherence of the gate within the high-dimensional space. The Mean Nearest-Neighbor (mNN) Purity Score serves as a quantitative metric of population density and cluster isolation. For each cell *i* within a gate *G*, we identify its *K*-nearest neighbors (*N*_*i, k*_) in the full ADT expression space. The purity for an individual cell is defined as:

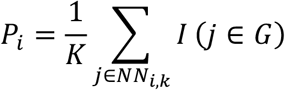

where *I* is an indicator function that equals 1 if the neighbor *j* is also within the gate *G*, and 0 otherwise. Results were benchmarked against a random baseline, defined as the null expectation of purity if cells were sampled at random from the total population (*N*_gated_/*N*_total_). Together, these results indicate that **scDIG**-defined gates are not artifacts of the two-dimensional embedding but correspond to coherent cell populations in the underlying high-dimensional ADT and RNA feature spaces.

## Discussion

### Comparison to other programs

Several recent tools have attempted to bridge the gap between automation and user control for CITE-seq data specifically. Single-Cell Virtual Cytometer introduces flow cytometry-like gating to multi-omics datasets, allowing users to create density plots based on antibody expression and perform lasso selections on 2D visualizations. However, this tool requires tab-separated text file inputs, limits data export to cell names from the final gated population only, and lacks integration with standard single-cell analysis workflows in R or Python. CITEViz, published in 2024, addresses some of these limitations by providing an R-Shiny application that accepts Seurat objects and enables sequential gating workflows similar to flow cytometry. While CITEViz successfully standardizes the gating process for CITE-seq data, it focuses primarily on surface protein markers and does not incorporate feature selection based on expression bimodality or density-based visualization of gene expression patterns in embedding spaces.

**scDIG** extends beyond these existing tools by combining several key innovations not available in a single platform. Unlike Single-Cell Virtual Cytometer or CITEViz, which focus on pre-defined protein markers, our tool implements Bimodal Index-based feature selection with adaptive sampling strategies, enabling systematic identification of genes and proteins with bimodal expression patterns across datasets of any size. To improve visualization of population boundaries, **scDIG** overlays contour lines derived from feature-weighted kernel density estimation onto embedding spaces, providing an objective, quantitative representation of cell density that complements traditional scatter plots.

Furthermore, **scDIG** operates directly on Seurat objects, maintaining compatibility with established scRNA-seq workflows.

### Limitations

**scDIG** is limited to Seurat objects (.rds) and cannot handle alternative formats such as SingleCellExperiment, AnnData, or flow cytometry files. Uploads are capped at 3 GB, and only one dataset can be analyzed per session.

## Availability of data and materials

The CAVA dataset analyzed during the current study is available available at GEO: GEO access number is GSE190570. The data will be made publicly accessible upon publication.

## Supplementary Material

**Figure S1.**
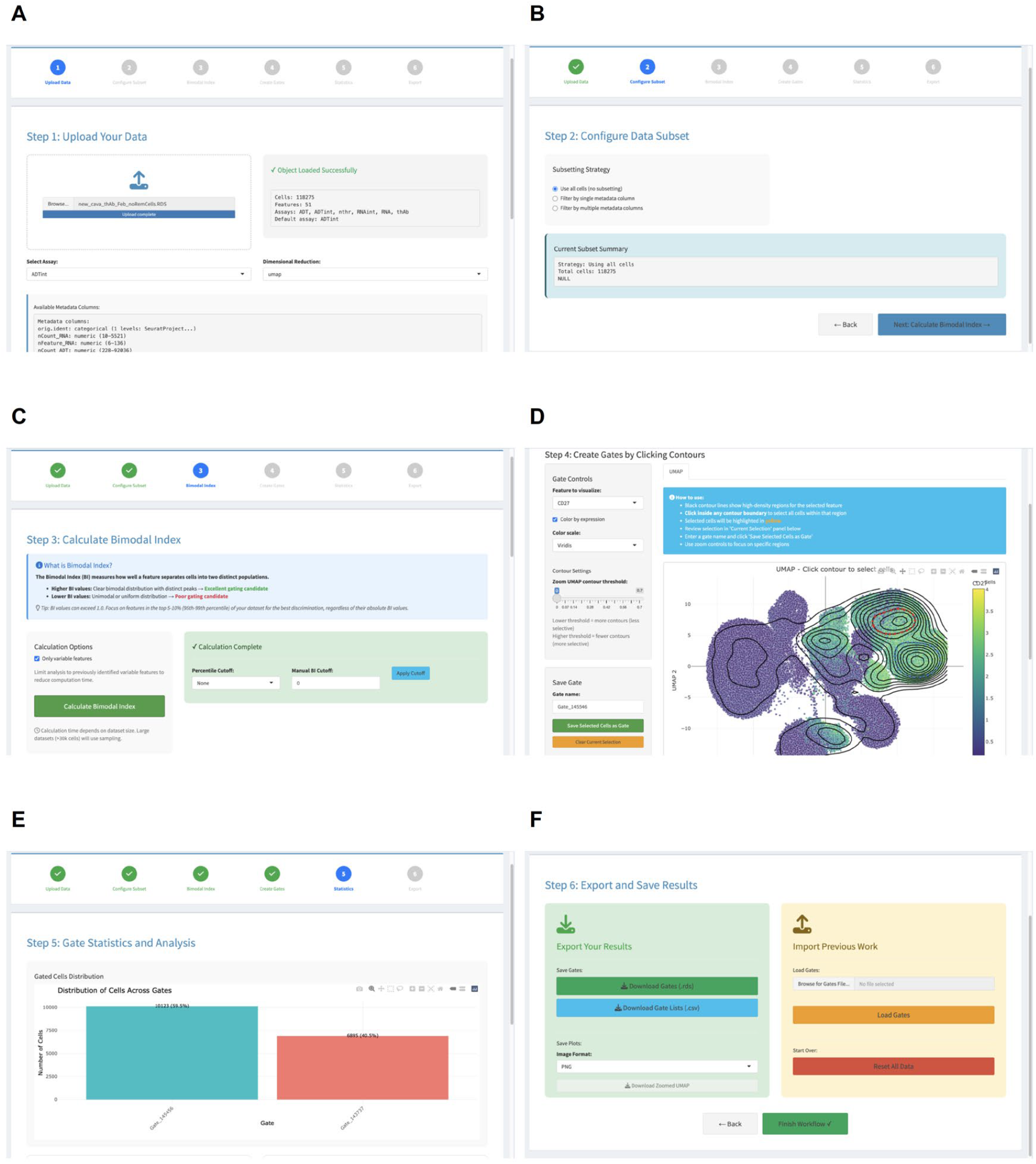
Screenshots of the scDIG application interface illustrating the six-step workflow. (A) Data upload and object validation. (B) Cell subsetting and filtering. (C) Bimodal Index calculation and feature selection. (D) KDE contour visualization and interactive gating. (E) Gate statistics and overlap analysis. (F) Gate export and import.

